# Assessing and removing the effect of unwanted technical variations in microbiome data

**DOI:** 10.1101/2021.05.21.445058

**Authors:** Muhamad Fachrul, Guillaume Méric, Michael Inouye, Sünje Johanna Pamp, Agus Salim

## Abstract

Varying technologies and experimental approaches used in microbiome studies often lead to irreproducible results due to unwanted technical variations. Such variations, often unaccounted for and of unknown source, may interfere with true biological signals, resulting in misleading biological conclusions. In this work, we aim to characterize the major sources of technical variations in microbiome data and demonstrate how a state-of-the art approach can minimize their impact on downstream analyses. We analyzed 184 pig faecal metagenomes encompassing 21 specific combinations of deliberately introduced factors of technical and biological variations. We identify several known experimental factors, specifically storage conditions and freeze-thaw cycles, as a likely major source of unwanted variation in metagenomes. We also observed that these unwanted technical variations do not affect taxa uniformly, with freezing samples affecting taxa of class *Bacteroidia* the most, for example. Additionally, we benchmarked the performance of a novel batch correcting tool used in this study, RUV-III-NB (https://github.com/limfuxing/ruvIIInb/), to other popular batch correction methods, including ComBat, ComBat-seq, RUVg, and RUVs. While RUV-III-NB performed consistently robustly across our sensitivity and specificity metrics, most other methods did not remove unwanted variations optimally, with RUVg even overcorrecting and removing some of the true biological signals from the samples. Our analyses suggests that a careful consideration of possible technical confounders is critical in the experimental design of microbiome studies to ensure accurate biological reading of microbial taxa of interest, and that the inclusion of technical replicates is necessary to efficiently remove unwanted variations computationally.

## Introduction

The technological advancement of sequencing technologies has made microbiome studies more accessible and meaningful. From amplifying short 16S rRNA hypervariable regions to taking advantage of long-read sequencing, the breadth of data options has enabled the field to flourish in the past couple of decades, allowing a better understanding of the role of microbiomes in numerous ecological, environmental and clinical contexts^1^. For example, the dynamics of the gut microbiome is now known to be influenced by environment and diet, and perturbations (or “dysbiosis”) have been linked to chronic conditions such as cardiometabolic diseases and type 2 diabetes (T2D)^2–4^. As a result, the human microbiome is now considered an important biological basis for potential therapeutic targets^5,6^. However, the multiplication of microbiome studies using a very wide range of technological methods and experimental designs comes with a cost on reproducibility. Any valuable application from microbiome studies has indeed been largely hindered by the lack of reproducibility due to the presence of unwanted technical variations.

Microbiome studies can differ considerably when it comes to the experimental approaches; and each step of the workflow, including variations introduced by the experimenter, has the potential to introduce artificial results due to unwanted technical variations^7,8^. For example, under-sampling might occur due to the lack of consideration during initial collection process, resulting in zero reads detected for certain microbiota that actually exist in the environment due to their underrepresentation in the samples^9^. Additionally, specific stool collection kits have been shown to impact microbial abundance of faecal samples differently^10^. DNA extraction, library preparation kits, storage condition, storage time, and choice of sequencing platforms have also been found to introduce artificial variations in microbial abundance^7,8,11,12^. As an example, outgrowth of certain microorganisms has been reported between sample preservation methods, as some methods are unable to prevent the growth of facultative anaerobic organisms^12^. Contamination of external microbial taxa can also contribute to unwanted variations and can happen at any stage of the experiment. For instance, recent studies have found that some library preparation kits used before sequencing could introduce specific microbial taxa coming from reagents in the kit^13^. This issue is observed to be even more critical in low-biomass samples, even leading to debated interpretations on the existence of a microbiome in environments that might not have any, such as the human placenta or the meconium^13–16^. Left unaddressed, such unwanted variations can considerably confound true biological signals and result in misleading conclusions.

Various computational methods have been developed to correct for batch effects within experiments. Popular methods such as ComBat^17^ and RUV^18^, originally developed for microarray and RNA-seq transcriptomics datasets, have also been considered for microbiome studies^7,19^ but do not particularly take into account the specific characteristics of microbiome data such as compositionality and zero-inflation. Indeed, contrary to features in transcriptomics datasets, the presence of each taxa is not independent from the rest of the taxa in the microbiome, and raw abundance information acquired after sequencing is not directly representative of the actual abundance in the environment due to the limit of the sequencing depth that each platform has^20^, a problem that is not commonly addressed in current studies^21^. Additionally, microbiome data are typically zero-inflated and very sparse, as many features are present in only a very few samples. Log-transformation of raw abundances, a process allowing for more robust statistical analyses to be performed, prerequires to substitute zero-values with a constant arbitrary number, also known as “pseudocount”. Despite being part of the compositional data analysis (CoDA) standards, there is an argument against adding pseudocount then using log-transformation in analyzing highly sparse datasets in the form of counts per sample, as it changes the ratio of taxa abundance substantially, diminishes variance from less abundant taxonomic groups and artificially exaggerates the differences between zero and non-zero values^22,23^.

In this study, we aim to characterize the major sources of technical variations in microbiome data and demonstrate how a state-of-the art approach can minimize their impact on downstream analyses. Using a dataset containing 184 faecal microbiome samples from pigs comprising up to 21 unique combinations of technical variations, we use RUV-III-NB (https://github.com/limfuxing/ruvIIInb/), a novel robust batch correction tool which utilizes Negative Binomial (NB) distribution to estimate and adjust for unwanted variations without the need to add pseudocounts, to identify parts of the experimental workflow that introduce critical unwanted variations that affect observed microbial abundances. Then, we compare the performance of RUV-III-NB to other popular tools including ComBat, ComBat-Seq, RUVg, and RUVs.

## Results

For the main analyses of assessing the contributions of unwanted variations towards microbial abundance, we utilized the novel RUV-III-NB, which is able to estimate and adjust sequencing count data for unwanted technical variations based on the negative control taxa and sample replicate information in the dataset. For benchmarking purposes, we refer to methods utilizing batch information for control measure (ComBat and ComBat-Seq) as ComBat-based methods, and methods utilizing control features (RUVg, RUVs, and RUV-III-NB) as RUV-based methods.

### Unwanted variations correlate with known sources of technical variations, and mostly affect highly abundant taxa

We acquired 7 unwanted factors (referred to as W1 to W7) from using RUV-III-NB on the spiked pig metagenomes, representing detected sources of unwanted variations within the dataset. The primary unwanted source of variation (called W1) correlated with log library size (r=0.648), which suggests that it is capturing the variability caused by sequencing depths between samples. It also correlated with log geometric mean (r=0.964), highlighting the ability of RUV-III-NB to identify the optimal form of normalization. In this case, it supports the method of utilizing log geometric mean as scaling factor, such as with the centered log-ratio (CLR) transformation, as a more effective approach than total sum scaling (TSS) normalization at providing the most accurate abundance estimate. A number of known experimental factors were also found to correlate well with other unwanted variations: we found storage conditions and freeze-thaw cycles correlating with W2 (r=0.770) and W3 (r=0.782), respectively **(Figure 1B)**. Differences in the library preparation kit used were also found to weakly correlate with W5 (r=-0.522) and W6 (r=0.500).

**Figure 1:**
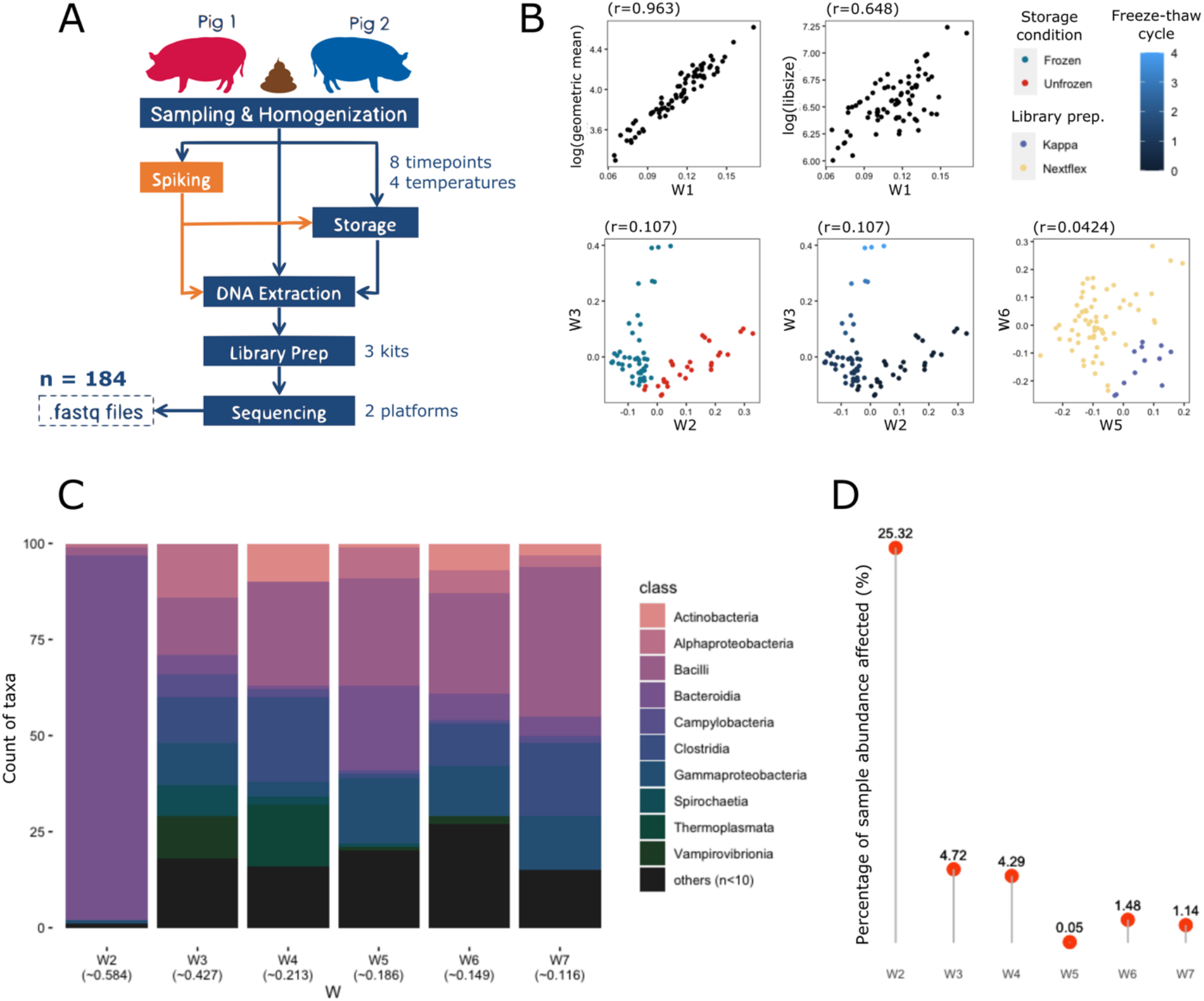
**(A)** Experimental workflow of the dataset, showing the sources of possible unwanted variations, some of which correlated with the calculated unwanted factors; **(B)** unwanted factor 1 (W1) correlated with log library size and log geometric mean, W2 correlated with storage conditions, W3 correlated with freeze-thaw cycles, and both W4 & W5 correlated with library preparation kits. **(C)** The top 100 microbial taxa most affected by each W colored by Class, ranked based on Veall-Zimmermann pseudo R-squared (mean listed under every W). Classes with less than two microbial taxa were merged as “others”. **(D)** The proportion of a sample’s total abundance affected by the top 100 microbial taxa for each W.

Microbial taxa belonging to the *Bacteroidia* class were found to be the most affected by W2, which correlated to storage conditions of the samples **(Figure 1C)**. Members of the *Bacilli* and *Clostridia* classes were also among the most affected across W3–W7. Incidentally, *Bacteroidia, Bacilli,* and *Clostridia* were the most abundant classes in the dataset, with 2,153 species belonging to *Bacteroidia,* totaling over 656 million reads (62.8% of total). To estimate the extent to which each of the detected unwanted factor affected our samples, we calculated the proportion of read counts belonging to the 100 most affected taxa, defined by the taxa with the highest pseudo-R^2^ for each unwanted factor (**Figure 1D**). We found that storage conditions (referring here whether samples were kept frozen or at room temperature before processing), represented by unwanted factor W2, affected up to 25.32% of a sample’s total abundance — a notable amount considering only 100 out of 8,453 taxa were taken into account **(Figure 1D)**. This suggests that after library size, storage conditions are the primary source of unwanted variation in microbiome studies. Unwanted factors W3 and W4 contributed to 4.72% and 4.29% of overall sample abundance, respectively, whereas W5–W7 each contributed to under 2%.

### Technical variations do not affect different taxa uniformly

The previous observation suggests that not every taxon is affected similarly by variations in technical procedures, and that some taxa are biologically more sensitive or resistant to varying experimental conditions than others. To further this, we analyzed abundances from several microbial classes in samples from the same source (P1) but subjected to different technical variations (**Figure 1A**). We found the highly abundant bacterial classes to still be differentially abundant between batch groups, including storage conditions, freeze-thaw cycles, and library preparation kits. Taxa from classes *Bacteroidia, Bacilli, Clostridia,* and *Alphaproteobacteria* were found to significantly differ in abundance when compared between storage conditions and freeze-thaw cycles, with taxa from *Alphaproteobacteria* also differed between samples generated from different library preparation kits **(Figure S1)**. Moreover, the differential abundance patterns, as in the state of the taxa in question being more abundant in one condition than the other, were not shared between bacterial classes in the same conditions: *Bacteroidia* was found to be more abundant in unfrozen samples, whereas *Bacilli, Clostridia,* and *Alphaproteobacteria* were less abundant. The pattern also differed for these classes in samples generated with varying freeze-thawing cycles and library preparation kits, with *Bacilli, Clostridia* sharing a similar abundance pattern across the different cycles. This suggests that the effect of technical variations is not uniform across different taxa.

### Most RUV-based methods remove unwanted variations better than ComBat-based methods

We then plotted relative log expression (RLE) plots before and after correction using ComBat, ComBat-seq, RUVg, RUVs, and RUV-III-NB, to assess how much overall unwanted variations were removed from each approach, by calculating a metric based on the variance of the medians and interquartile ranges. Unlike with factors of unwanted variations or principal components (PCs) where we could assess how each of them correlates with specific technical variations, RLE plots only assess the whole/overall unwanted variations that exist within a dataset; hence, the term overall unwanted variations here is used when referring to results from RLE analysis. The RLE plots and principal component analysis (PCA) plots before and after RUV-III-NB correction (**Figure 2A, Figure 2B)** visually represent the removal of overall unwanted variations. Based on the RLE metric, better corrections were seen for RUV-based methods than ComBat-based methods, with the exception of RUVs using spike-in taxa only **(Figure 2C)**. RUVg had the best RLE metric in average 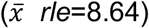, followed by RUV-III-NB 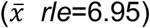. RUVs removed unwanted variations comparably to RUV-III-NB when using both combination and only empirical sets of control features yet did not remove much overall unwanted variations when using only spike-in taxa (*rle*=0.49), scoring even lower than uncorrected data (log2 *rle*=2.93, CLR *rle*=3.98). Both tested ComBat methods, using log2 and CLR transformations, had RLE metrics higher than uncorrected data, but still scored lower on average 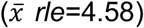 than the lowest-scoring RUVg (*rle*=5.15) and RUV-III-NB (*rle*=4.98) runs. ComBat-Seq did not seem to remove the entirety of the unwanted variations as the resulting RLE metric was similar to uncorrected data with log2 transformation (*rle*=2.94). This analysis suggests that RUV-based methods generally perform better in removing overall unwanted variations in microbiome data than ComBat-based methods. To get a visual representation of overall variation in datasets and the effects of correcting for batch effects, we performed principal component analysis (PCA) on the microbiome composition matrix data and calculated the silhouette statistics (ss) for clustering by a technical factor of main batch separation (in this case storage conditions), using the top four principal components (PCs) before and after correction (**Figure 2B**). Specific clustering of samples based on storage conditions and the library preparation kit used could be seen in the PCA plots before any batch effect correction, specifically captured by PC3 and PC4, respectively, but this was no longer apparent after correction with RUV-III-NB using both spike-in and empirical taxa **(Figure 2B)**. With the exception of RUVs using only empirical control taxa *(ss*=0.51), all approaches had lower silhouette scores compared to uncorrected data (uncorrected *ss*=0.488; ComBat *ss*=0.11-0.26; ComBat-Seq *ss*=0.11; RUVg *ss*=0.14-0.17; RUV-III-NB *ss*=0.09-0.16; RUVs *ss*=0.08-0.51), suggesting successful removal of storage condition effects **(Figure S2)**. The effect of using different normalization methods did not seem to alter the silhouette scores in uncorrected data, unlike when using ComBat approaches. Overall, correction using RUV-III-NB and ComBat-Seq yielded the lowest average scores within this dataset, indicating their robustness in removing unwanted variations from the major technical variation source.

**Figure 2:**
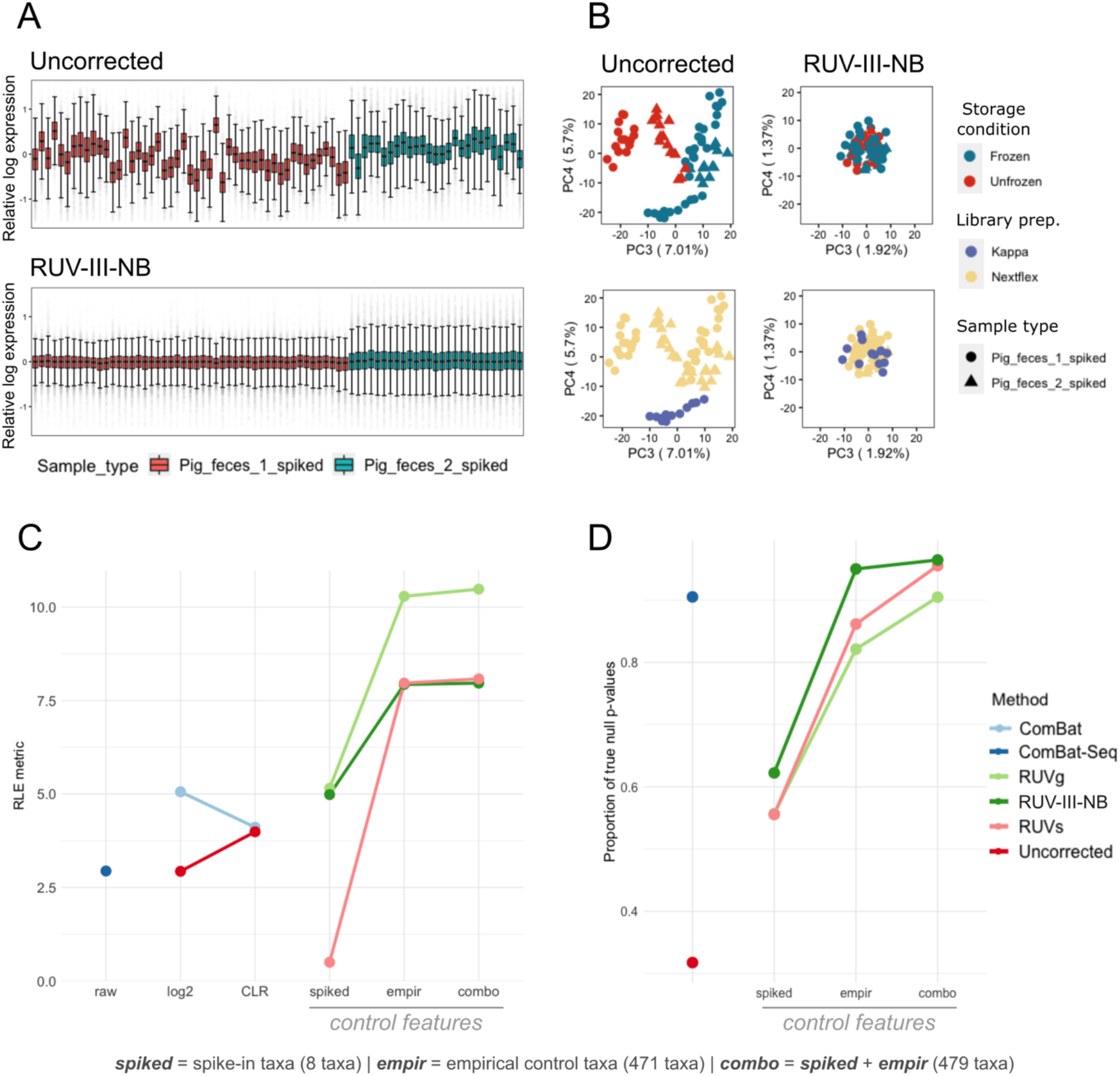
**(A)** RLE plots of all spiked samples, before correction (top) and after RUV-III-NB correction with combination control taxa (bottom). In RLE plots, successful removal of unwanted variations is represented by the medians of sample from the same individual being as close to zero as possible and as linear to each other as possible, as well as similar interquartile ranges (IQR) between the samples visualized by the size of the boxplot. **(B)** PCA plots of the same set of samples, before (left) and after RUV-III-NB correction with combination control taxa (right). Clustering based on storage conditions and library preparation kits could be explained prior to correction by PC3 and PC4, respectively – which is no longer the case after correction. The lack of visible clustering suggests successful removal of unwanted variations. **(C)** Comparison of correction method performances based on relative log expression (RLE metric), where higher number indicates better removal of overall unwanted variations. With the exception of RUVs with solely spike-in taxa as control features, RUV methods in average performed better compared than ComBat-based methods. **(D)** Proportion of true null p-values (pi0) of uncorrected and corrected data using different correction methods after differential abundance analysis. Since storage conditions were found to be the main batch variable in this dataset, comparison was done between frozen and unfrozen samples of samples from the same source (pig 1). Since edgeR requires integer counts as input, ComBat was omitted from this comparison. RUV-III-NB resulted in the highest pi0 overall when using the combination set of control taxa, and still performed better than ComBat-seq when using solely empirical taxa as control features.

To identify taxa most affected by different unwanted sources of variation, we performed differential abundance analysis using edgeR between the different storage conditions, in which only spiked P1 samples were used. After FDR correction and fold change filtering (FDR<0.05, ļlog2(FC)ļ >1), samples ran using ComBat-Seq had the lowest number of differentially abundant taxa after correction, followed by RUV-III-NB and RUVs with combination control taxa **(Table S1)**. Yet, correction with RUV-III-NB resulted in the highest estimates of the proportion of null (pi0; non-differentially expressed) features, with correction using combination and solely empirical taxa reaching 0.964 and 0.950, respectively — higher than ComBat-Seq (pi0=0.905) **(Figure 2D)**. Correction using RUV-III-NB also resulted in the highest average pi0 compared to the other RUV methods, followed by RUVs which performed comparably when using combination control features (pi0=0.955). RUV methods with solely spike-in control taxa all performed comparably, with higher results than uncorrected data but lower than ComBat-Seq. Taken together, these observations suggest that RUV-III-NB performed the most consistently robust compared to the other methods in removing unwanted variations.

### Technical replicates and choice of control features are crucial for RUV methods

To assess how well correction methods preserve individual information, we calculated silhouette statistics of individual separation after removal of unwanted variations, a metric describing how well samples from two conditions (here, pigs P1 and P2) separate after clustering, based on how close each point in one cluster is to points in the neighboring clusters; in this case, high value indicates precise clustering and separation of samples based on the pig individuals and, whereas low value suggests otherwise. RUVs using empirical control taxa had the highest silhouette score (ss=0.797), followed by RUVs and RUV-III-NB with combination control taxa (ss=0.790 and 0.761, respectively) **(Figure 3A)**. However, RUV-III-NB in average had a higher silhouette score than RUVs, as correction with solely spike-in control taxa using the latter yielded a silhouette score lower than uncorrected data (RUVs 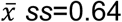, RUV-III-NB 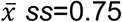). ComBat-based methods performed comparably to uncorrected data, with correction using ComBat using log2 having higher silhouette score. Interestingly, RUVg yielded the lowest score overall, with all three control feature options scoring just below zero. This is visualized in the PCA plots before and after correction, in which separation of P1 and P2 samples are explained by PC1 and PC2 even before any correction. The discrimination between individual metagenomic profiles is retained after RUV-III-NB correction, but not after using RUVg **(Figure 3B)**. This analysis suggests that sample replicate information is necessary when correcting unwanted variations using RUV-based methods to avoid over-correction. That being said, despite both methods utilizing sample replicate information, RUV-III-NB still prevailed over RUVs when taking into consideration performance consistency over the different sensitivity and specificity metrics.

**Figure 3:**
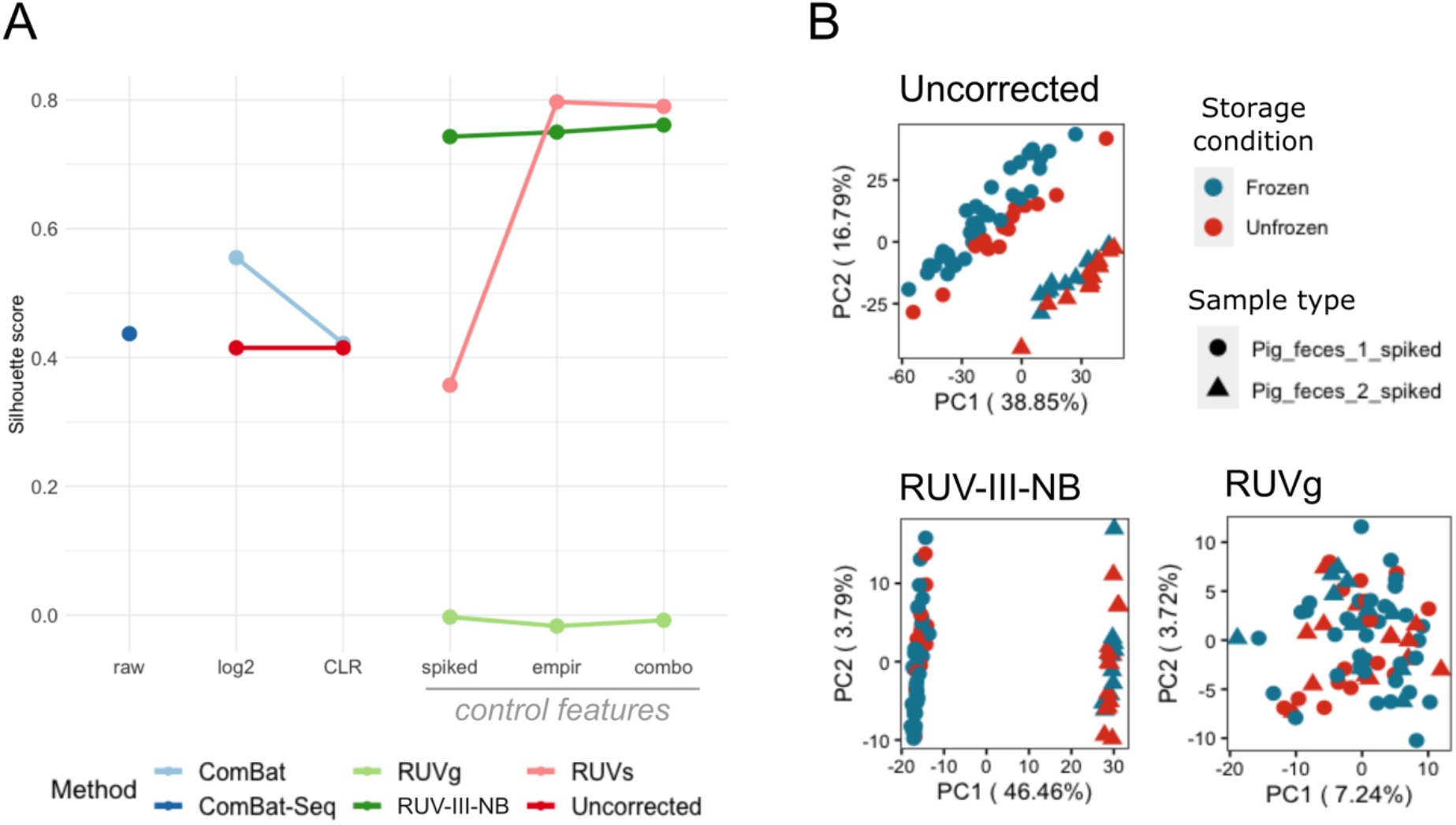
**(A)** Comparison of correction method performances in preserving individual biological information through silhouette scores, which are calculated based on the main principal components (PCs) of spiked samples and explain how well samples separate between defined groups (higher = better). RUVs using solely empirical control features had the highest silhouette score in separating pig 1 and pig 2 samples, yet correction using RUV-III-NB had the highest average silhouette score overall – indicating its consistency despite the different set of control features. RUVg performed really poorly overall, with silhouette scores using all the different control features placing significantly lower than even uncorrected data. **(B)** Based on the PCA plots of all spiked samples, separation of pig 1 and pig 2 samples can be explained by the main PCs in uncorrected data (top) and is even further defined after RUV-III-NB correction, clearly explained by PC1 (bottom-left). Separation of the two pig sources is lost after RUVg correction, as the two defined clusters are no longer visible.

## Discussion

The impact of technical variations in microbiome analysis is an important topic that has been explored numerous times, and past studies have reported on the best practices at every stage of sample processing^24^. However, our study is the first to quantify the contribution of different technical factors towards unwanted variations in a shotgun metagenomic sequencing dataset, and how to best remove them. To do so, we used RUV-III-NB, a novel tool accessible on (https://github.com/limfuxing/ruvIIInb/), which we also compare to other available published methods.

### Inconsistent sample storage can be a large source of unwanted variations in microbiome studies

The variety of storage conditions – which encompass factors such as temperature, time, and storing method used – have been previously found to introduce considerable effects on microbial abundance^11,12,25,26^. The impact of different storage conditions is also not limited to metagenomic data, as Hickl et al. also reported significant differences in microbial protein identification between samples stored using flash freezing and RNAlater^27^. Our study does not aim to replicate and validate the findings of such studies, but to show how, if left unaddressed, the differences in sample storage conditions could introduce significant changes in microbial abundance of abundant taxa which are potentially important for our understanding of particular conditions or diseases. The three most-affected bacterial classes in our study, namely *Bacteroidia*, *Clostridia* and *Bacilli,* are among the most dominant microbial classes in the mammalian gut^28–30^, therefore it is not surprising to see them strongly affected by technical variations. Both *Bacteroidia* and *Clostridia* classes in particular have been found to be significantly impacted by different storage methods in both metagenomic and metaproteomic data^27^. In clinical cases, the dysbiosis of members of the classes *Bacteroidia* and *Clostridia* have been linked to a wide variety of conditions, including T2D, Crohn’s disease, lupus, HIV infection, and major depressive disorders^30–35^. Hence, extra precautions are necessary in making sure the experimental design minimizes sources of unwanted variations as possible.

### Metagenomic sequencing depth and data normalization also contribute to unwanted variations in microbiome studies

Aside from storage conditions, RUV-III-NB identified that library size and geometric mean correlated with unwanted factor W1 in our study, which is unsurprising. In microbial metagenomics, sequencing depth typically impacts the robust classification of reads into genes or microbial taxa, and optimal depth to use depends on each study objectives^36,37^. Our dataset included a range of sequencing depths, which explains why library size was captured as a source of variation by RUV-III-NB. Furthermore, although normalization and log transformation alone are not enough to remove unwanted variations, normalization by library size, also known as TSS normalization, remains widely used in the field mostly due to conceptual simplicity^38^. However, this approach does not account for heteroskedasticity and is very sensitive to pseudocount^39^. With unwanted factor W1 correlating much stronger with geometric mean than library size, our work suggests that normalizing by geometric mean (e.g., CLR transformation) rather than by library size (e.g. TSS) potentially limits unwanted variations in observed microbial abundances.

### RUV-III-NB is a robust and consistent microbiome batch correction method

Although the observation of best practice methods and preventive measures are always recommended when it comes to avoiding unwanted technical variations in any experimental setting, there are a few existing computational approaches to rectify batch effect variation after a dataset has been processed and sequenced. Here, we show that, despite deliberately introduced complications in our gut metagenomics experimental workflow, the effective removal of unwanted variations was still possible during downstream analysis, with different methods varying in efficiency. Methods such as ComBat, RUVg, and RUVs require normalization of the data prior to the actual batch removal step, but RUV-III-NB is able to take into account the most optimal normalization method and identify unwanted variations in one step without any prior pseudocount addition.

Additionally, ComBat-based methods rely on known batch information and might not be suitable in the realistic situation of multiple batch effects conflating with true biological signals. Nevertheless, when batch information is clearly defined, ComBat-based methods remove unwanted variations efficiently from our samples, but also impact the recovery of true biological signals separating individuals. In our work, ComBat-based methods were also unable to identify and adjust for unknown unwanted variations, which are expected to be common and unavoidable in reality. RUV methods avoid this hurdle by adjusting samples based on the variations of the control features, which are assumed to be in constant amount across technical variations.

One potential issue with RUV methods is that, unlike with RNA-seq data where housekeeping genes have been established^40,41^ and are routinely used to normalize expression, there is no obvious parallel in microbiome features, as taxa composition varies considerably across datasets. The choice of control features is especially crucial for the methods to work as intended; therefore, the presence of spike-ins is highly beneficial when utilizing RUV methods in microbiome studies. In our case, the mock community consisted of 6 bacterial and 2 eukaryotic microorganisms. Over 3,000 genes have been established to be consistently expressed in human cells and are used to normalize expression in RNA-seq studies, which is considerably larger than our spike-ins. We therefore supplemented the spike-in taxa with a set of empirical control taxa. In this study, correction with only the spike-in control taxa removed unwanted variations to an extent, though using the additional empirical control taxa significantly improved it. Additionally, we observed that using only empirical taxa without spike-ins was still effective in removing unwanted variations, indicating that it is possible to effectively use RUV methods without spike-in taxa, which is more convenient when preparing samples.

When comparing between RUV methods, we showed how inclusion of sample replicate information is important, as demonstrated by RUVg’s tendency to overcorrect and lose individual separation in the process. Both RUVs and RUV-III-NB utilize sample replicate information, and are therefore able to avoid overcorrection, yet the former performed less consistent across the performance tests. RUV-III-NB has an advantage over RUVs due to its direct modelling of meanvariance relationship of count data – a feature not shared by RUVs.

In this study, we show how technical variations – especially storage conditions – may introduce unwanted variations in microbiome data, affecting the observed abundances of important and dominant microbial taxa. Left unaddressed, this might be problematic for studies focusing on such taxa, therefore careful considerations are crucial in designing the experimental workflow of microbiome studies. Minimizing the possibility of introducing unwanted variations by limiting the presence of batches and utilizing consistent storage conditions and equipment are highly suggested. We finally show that for existing datasets, post-processing corrective measures can still be performed *in silico* to remove unwanted variations stemming from variations in experimental techniques, and we suggest the use of *RUV-III-NB* as a consistent and robust method.

## Materials and Methods

### Sample processing, library preparation, and sequencing conditions

The fecal samples investigated in this study originated from two individual pigs (P1 and P2) collected right after defecation from two different conventional pig farms in Denmark as previously described^42,43^. Each individual pig faecal sample was homogenized with a sterile wooden spatula and then separated into two large aliquots. One of the aliquots was spiked with a freshly prepared mock community consisting of 6 bacterial and 2 eukaryotic microorganisms, namely *Propionibacterium freudenreichii* DSM 20271, *Bacteroides fragilis* NCTC 9343, *Staphylococcus aureus* NCTC 8325, *Fusobacterium nucleatum* ATCC 25586, *Escherichia coli* ATCC 25922, *Salmonella* Typhimurium str. ATCC 14028S, *Cryptosporidium parvum* IOWA II isolate, and *Saccharomyces cerevisiae* S288C. The spiked fecal samples were homogenized using a sterile wooden spatula, and small aliquots for each sample storage condition were prepared in Eppendorf tubes for both, the spiked and unspiked aliquots. DNA was isolated immediately from the aliquots for the initial time point (storage for 0h) using a modified QIAamp Fast DNA Stool Mini Kit (Qiagen) protocol with an initial bead beating step (MoBio garnet beads)^44^. The remaining aliquots were stored at different storage conditions comprising different temperature and time combinations **(Figure 1A)**. Samples were stored at 22°C, 5°C, −20°C, and −80°C for several hours (days) (16h (0.67 days), 40h (1.6 days), 64h (2.6 days), 88h (3.6 days)) and for the temperatures −20°C and −80°C also for months and up to one year (4m, 8m, 12m). Aliquots stored for 40h and 88h, as well as a subset of 64h, also underwent 2–4 freeze-thaw cycles. All fecal samples underwent the same DNA isolation method (see above)^43,44^ prior to library preparation.

Three different library preparation kits (the PCR-free NEXTflex and KAPA, as well as Nextera) and two sequencing platforms (HiSeq 4000 and NextSeq 500) were used in the study, similarly as previously described^42,43^. All samples were sequenced paired-end with a read length of 150 bp. Two to three technical replicates were performed for each treatment combination.

Together, a total of 184 different samples spread across 60 different replicate groups were acquired, encompassing 21 specific combinations of storage conditions, spiking status, library preparation kit used, as well as thawing cycles. The raw reads are deposited at the European Nucleotide Archive (ENA) (Project acc.: PRJEB31650).

### Quality control and taxonomic classification

Quality control of sequencing files was done following DTU Food’s in-house pipeline, FoodQCpipeline (https://bitbucket.org/RolfKaas/foodqcpipeline), in which BBMap’s bbduk2 (v38.71, https://jgi.doe.gov/data-and-tools/bbtools/) was used to trim reads with a length of at least 50bp, Phred score of at least 20, and also remove a custom list of Illumina adapters. FastQC (v0.11.8) was also applied to the files before and after trimming to assess the quality of the reads^45^.

Taxonomic classification was done with Kraken2^46^, followed by taxa abundance re-estimation using bracken^47^ at species level with read length of 150 and minimum taxa threshold of 1. A custom genomic index was used for the classification based on GTDB release 89^48,49^, which includes 23,458 bacterial and 1,248 archaeal species. After merging individual reports, taxa filtering was done based on CPM>4 in at least 15% of all samples. A total of 8,453 taxa were included for downstream analysis.

### RUV-III-NB batch correction method

The novel RUV-III-NB (https://github.com/limfuxing/ruvIIInb/) method extends the RUV-III method that was previously developed for array-based gene expression data^50^. Instead of using linear model, it uses a Negative Binomial-based generalized linear model (GLM) with log link function to model the effect of wanted biological signals and unwanted variations on the sequencing count at taxon-level. The method uses negative control taxa (i.e., taxa whose variations are assumed solely due to unwanted variations) to estimate the latent unwanted factors, followed by the utilization of replicate information to estimate the taxon-specific effect of these latent unwanted factors on the variations of the sequencing count. Once these are estimated, RUV-III-NB uses percentile-invariant adjusted count to output the sequencing count matrix that has been adjusted or corrected for the unwanted variations. The percentile-invariant adjusted count was calculated based on a modification version of the randomized quantile residual method^51^.

### Sensitivity and specificity of batch correction methods

We compare the effectiveness of the following methods for removing unwanted variation: ComBat^52^, ComBat-Seq^53^, RUVg^18^, RUVs^18^, and RUV-III-NB (see RUV-III-NB subsection). ComBat and ComBat-Seq both utilize known batch variable to remove batch effects from a dataset, though the former allows normalized and/or log-transformed matrix as input, whereas the latter requires raw integer count matrix as input^52,53^. RUVg, RUVs, and RUV-III-NB all utilize control features in estimating unwanted factors. Both RUVg and RUVs assume an underlying Normal model and take normalized and/or log-transformed matrix as input, while RUV-III-NB only accepts integer count matrix; though both RUVs and RUV-III-NB require sample replicate information^18^. Out of all the methods, only ComBat-Seq and RUV-III-NB deal directly with the mean-variance relationship in the count data by using NB distribution. We tested two different log-transformation approaches for uncorrected data and prior to ComBat correction: normalization by library size / total sum scaling (TSS) followed by log2 transformation, as well as Aitchison’s centered log-ratio (CLR) transformation^54–56^. Prior to RUVg and RUVs, CLR transformation was also performed. For RUV methods, we set *k* – which represents the number of unwanted variations – to 7 as it is the highest possible number for the smallest set of control features. For RUV-III-NB, we also set the parameters *lambda.a=0.01* and *lambda.b=5.*

In addition to the 8 spike-in taxa, we curated a set of empirical control taxa – which is defined as a set of microbial taxa that are present and empirically concordant in abundance in both groups that we would like to compare against – to be used as control features in RUV methods. To identify the empirical control taxa, the following procedure was used:

- Differential abundance analysis was done between spiked and unspiked samples using edgeR^57,58^ for each unique replicate group of each pig. In average, only <1% of the taxa were significantly differentially abundant in all the replicate groups.
- We then took the intersection of the least significant taxa from each replicate group within the same pig, specifically the bottom 86% of overall taxa based on FDR<0.05, as control taxa for each pig.
- From the control taxa of both pigs, we then took the intersection and identified a total of 471 overlapping taxa. This set is what we refer to as the empirical control taxa from this point on.

For each RUV method, performance was assessed while using either solely spike-in taxa (8 taxa), solely empirical taxa (471 taxa), or the combination of both as control features (479 taxa).

We generated relative log expression (RLE)^59^ and principal component analysis (PCA) plots before and after correction to assess how well the different methods perform. RLE plots visualize the presence of unwanted variations by calculating the deviations of from the median of each feature, in this case microbial taxa, whereas PCA plots visualize clustering of certain batches through unsupervised dimensionality reduction^59,60^. For RUV-III-NB, the log of percentile-invariant adjusted count was used as the adjusted data matrix for the visualizations. To compare the RLE plots between methods, we calculated a metric capturing the variance of medians of samples from the same individual as well as the variance of the RLE’s interquartile ranges (IQR):

- We subtracted each sample’s median with the average median of all samples from the same pig source, then find the variance of all the medians.
- After the addition of the variance of medians and the variance of all the IQRs from the RLE, we then performed log transformation to the total.
- We took the negation of the transformed value; hence, higher metric implies more effective removal of unwanted variations.

For PCA plots, we calculated the silhouette statistics using the ‘cluster’ package ^61^ (version 2.1.0) to assess two different comparisons:

- Preservation of individual source information, in which we calculated the score of separation between P1 and P2 samples and higher scores signify better preservation.
- Batch dispersion, in which we calculated the scores to see clustering of batch groups within each pig group, with lower scores signifying better removal.

We also performed an additional specificity assessment for the methods through differential abundance analysis using edgeR^57,58^ on samples from the same source (P1) between groups of the main batch – in this case, between frozen and unfrozen storage conditions. Since ComBat only produces a transformed adjusted data matrix and not the estimated factors of unwanted variations, and therefore lacking the factors of unwanted variations necessary for covariates in differential abundance analysis, it is excluded from this analysis. Output from ComBat-Seq was directly used as input for the analysis. For RUV methods, untransformed integer counts were used as input along with factors of unwanted variations (W) as covariates in the linear model. Out of all the differentially abundant microbial taxa, we set the significance threshold to those with p-value < 0.05 after FDR correction and with absolute log-foldchange of over 1. We also performed q-value estimation of the results, which measures the proportion of true null p-values (pi0).

### Quantify relative contribution from unwanted factors

We used RUV-III-NB to quantify unwanted factors in the data and analyze their relative contribution towards microbial taxa abundance. Hence, for our main analysis only spiked samples were used as the approach requires control features as input, which in our case included the 8 spike-in taxa and an additional set of 471 empirically constant taxa explained in the previous section. We set the same number of *k* as stated in the previous section for consistency. We then used a negative binomial generalized linear model *(glm.nb)* and model the regression of each microbial taxa as the response variable and the progressive accumulation of factors of unwanted variations W as the predictor. We then calculated the Pseudo-R^2^ from each model using the PseudoR2 function from the ‘DescTools’ R package (version 0.99.38)^62^ with Veall-Zimmermann correction^63^, as it is among the closest approximations to ordinary least square R^2^. Excluding the first factor, the contribution of each individual unwanted factor (W) towards each microbial taxon is represented by the difference in pseudo-R^2^ between the cumulative n^th^ and n-1^th^ models (i.e. contribution of W3 was calculated by subtracting the pseudo-R^2^ accounting for unwanted factors W1+W2+W3 with the value accounting for unwanted factors W1+W2). To see the effect of known technical variations on microbial abundance, we then took the top 100 affected microbial taxa by each of the individual factor of unwanted variations, ranked based on their pseudo-R^2^ and grouped based on taxonomic class. Since factor of unwanted variation W1 correlated strongly with library size and geometric mean instead of any known technical variation, we only included factors W2 – W7 in this step. From the top 100 affected microbial taxa in each factor of unwanted variation, we calculated the proportion of their abundance in an average sample. For each of the most dominantly affected microbial classes, we also performed Wilcoxon signed rank test between storage conditions and Kruskal-Wallis test to compare the different freeze-thawing cycles and library preparation kits on log2-transformed, TSS-normalized data.

## Authors’ contributions

AS, MI, and MF conceived the original project. AS and GM supervised the experiments. AS constructed the tool highlighted in the study. SJP provided the microbiome dataset used in the study. MF performed the experiments, analyzed the data, prepared the figures, and wrote the bulk of the manuscript. GM, MI, AS, and SJP wrote and revised the manuscript. All authors read and approved the manuscript.

## Acknowledgements

MI was supported by the Munz Chair of Cardiovascular Prediction and Prevention. MF was supported by a Melbourne Research Scholarship from The University of Melbourne jointly funded by the Baker Heart and Diabetes Institute. This work was supported by Health Data Research UK, which is funded by the UK Medical Research Council, Engineering and Physical Sciences Research Council, Economic and Social Research Council, Department of Health and Social Care (England), Chief Scientist Office of the Scottish Government Health and Social Care Directorates, Health and Social Care Research and Development Division (Welsh Government), Public Health Agency (Northern Ireland), British Heart Foundation and Wellcome. This study was also supported by the Victorian Government’s Operational Infrastructure Support (OIS) program.

The funders had no role in study design, data collection and analysis, decision to publish, or preparation of the manuscript. The views expressed in this manuscript are those of the author(s) and not necessarily those of the NIHR or the Department of Health and Social Care.

## Data Availability

The raw sequence data was deposited at the European Nucleotide Archive (ENA) under accession number PRJEB31650.

## Supplementary Materials

### Supplementary Table

**Table S1:**
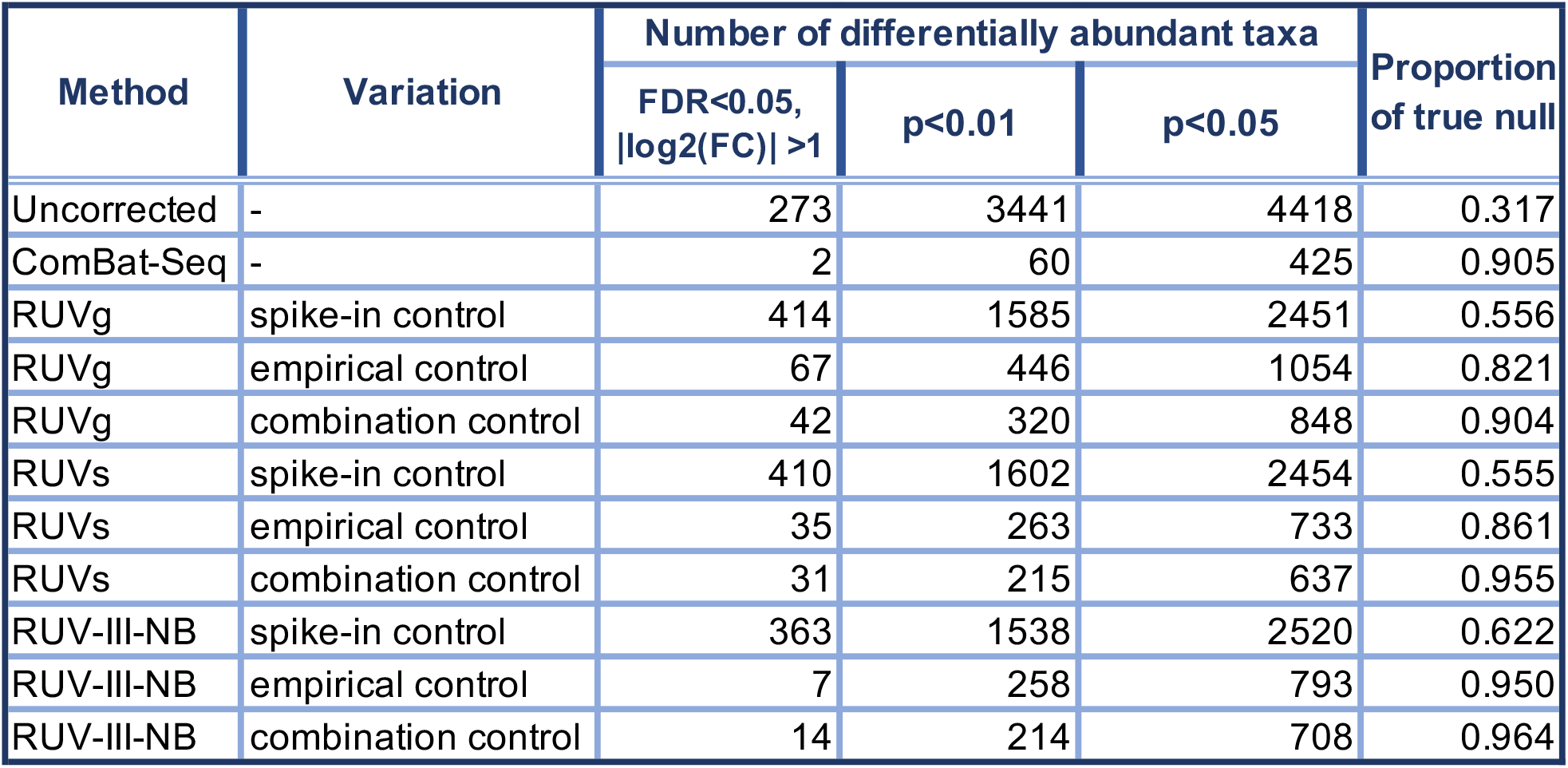
Results of differential abundance analysis of spiked pig 1 samples between storage conditions (frozen vs unfrozen) using edgeR. Since all originated from the same individual and were spiked, little-to-no taxa should be detected as differentially abundant. Out of 8,355 taxa, our analysis with ComBat-Seq detected the lowest number of differentially abundant taxa (2) in the dataset, yet RUV-III-NB with combination control taxa resulted in the highest proportion of true null (pi0), indicating higher specificity.

### Supplementary Figures

**Figure S1:**
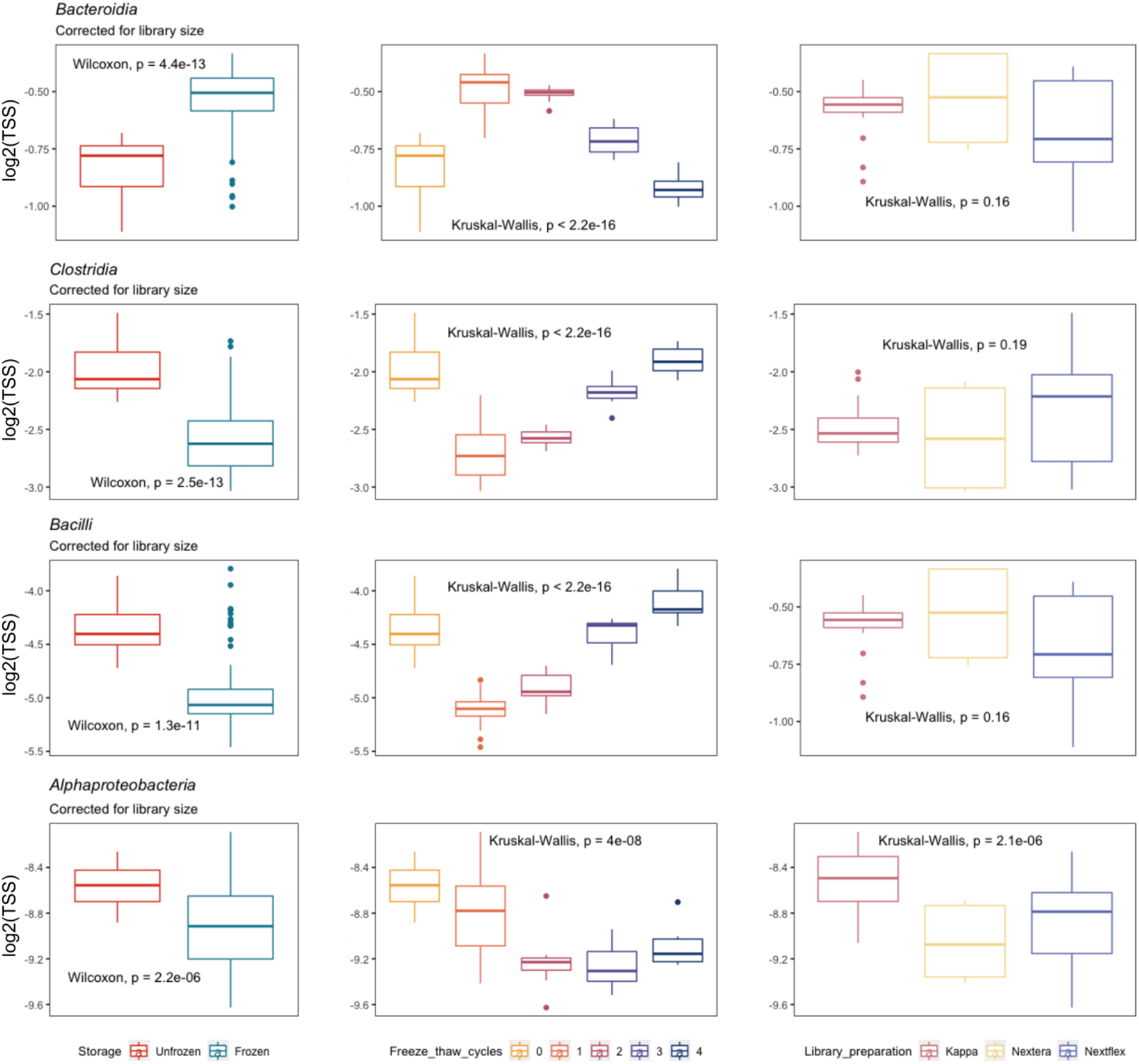
Differential abundance between variables of different batches (storage conditions, freeze-thaw cycles, and library preparation kits) in four of the most dominantly affected bacterial classes by unwanted variations: Bacteroidia, Clostridia, Bacilli, and Alphaproteobacteria. All classes were found to be differentially abundant between storage conditions and freeze-thaw cycles, with Alphaproteobacteria also being differentially abundant between library preparation kits.

**Figure S2:**
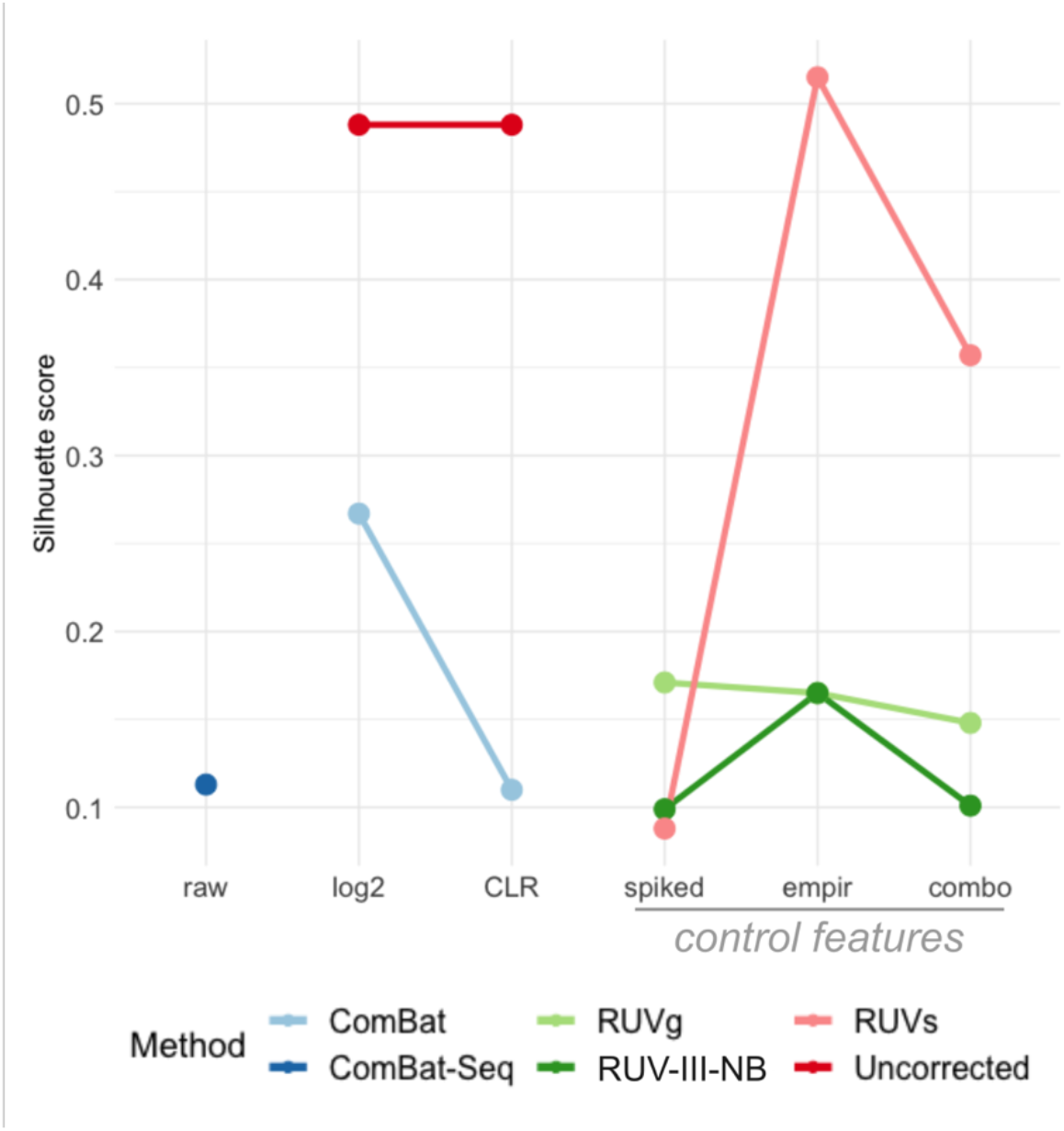
Comparison of correction method performances in removing unwanted variations from the main source (storage conditions). We calculated silhouette statistics based on the main principal components (PCs) of spiked samples, in which lower scores indicate poor clustering of batch variables, and therefore better removal of unwanted variations. With the exception of RUVs using only empirical control taxa (ss=0.51), all approaches had lower silhouette scores compared to uncorrected data (uncorrected ss=0.488; ComBat ss=0.11–0.26; ComBat-Seq ss=0.11; RUVg ss=0.14–0.17; RUV-III-NB ss=0.09–0.16; RUVs ss=0.08-0.51).

**Figure S3:**
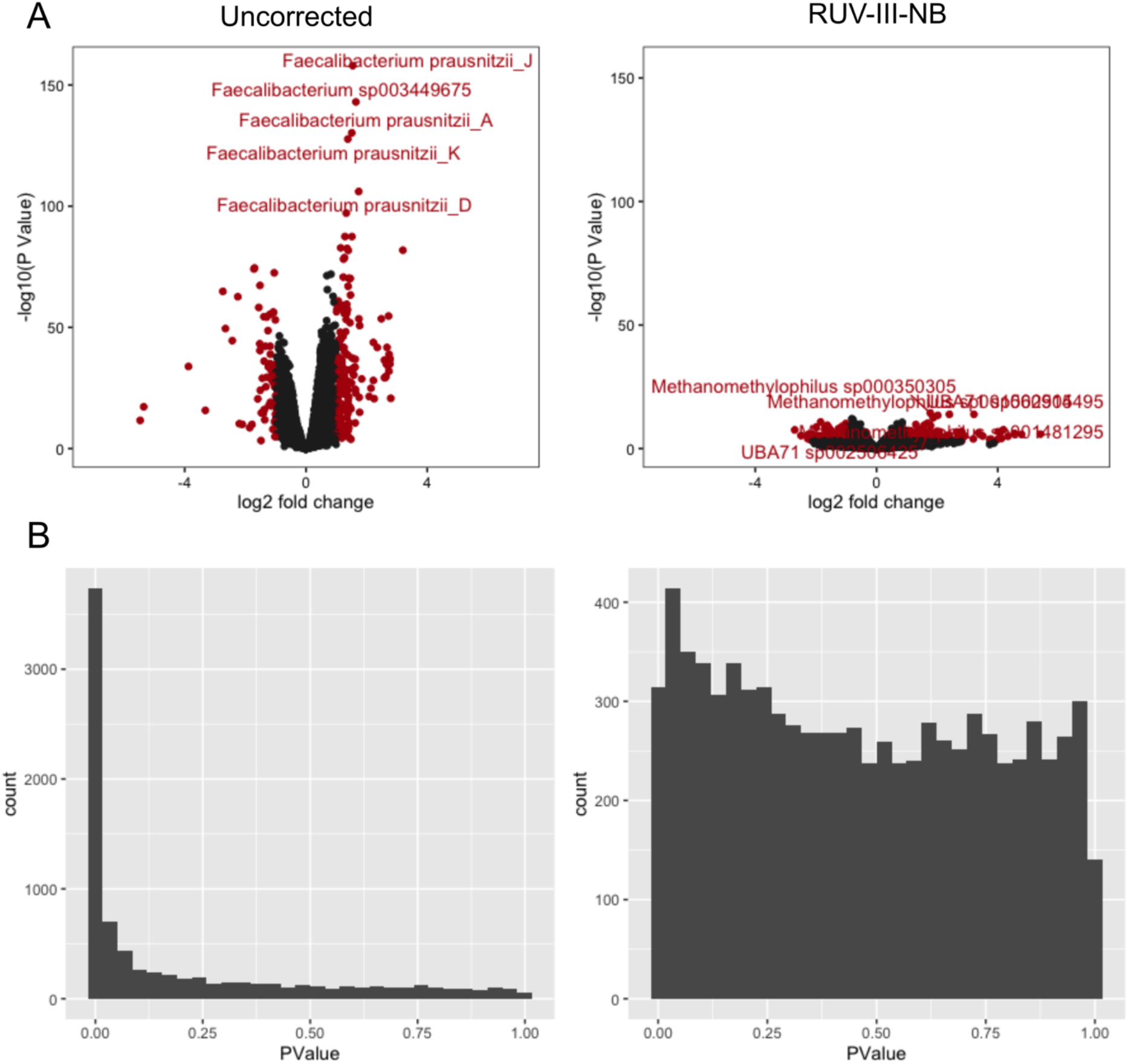
Results of differential abundance analysis of spiked pig 1 samples between storage conditions (frozen vs unfrozen), before (left) and after (right) correction using RUV-III-NB (with combination control features). Since the samples come from the same biological source, there should be little-to-no differences between the storage conditions. **(A)** volcano plot before correction shows over 200 significantly differentially abundant taxa (FDR<0.05, |log2(FC)|>1), with only 14 differentially abundant taxa after RUV-III-NB correction. **(B)** The distribution of p-values before correction is anti-conservative, noted by the peak on the left due to all the differentially abundant taxa. A more uniform p-value distribution can be seen after RUV-III-NB correction, suggesting minimum number of differentially abundant taxa.

